# Frontal β bursts segregate into rhythmically distinct regimes that differentiate cognitive states

**DOI:** 10.1101/2023.11.13.566594

**Authors:** Zachary D Langford, Emmanuel Procyk, Charles R E Wilson

## Abstract

Beta band rhythms often appear as brief bursts, but how variations in burst properties impact neural function is unclear. We probed beta burst heterogeneity by developing two complementary detection algorithms. One isolates brief high amplitude events (bursts of power, BoP) and another that identifies consistent oscillations that span multiple cycles (bursts of consistency, BoC). Examining frontal LFP and ECoG recordings from mice and macaques, these two burst types occupied the same 15 to 30 Hz frequency band yet showed minimal temporal overlap, indicating independent phenomena probably with distinct neural generators. Crucially, when task demands shifted between high and low cognitive control states, BoC bursts were enriched during demanding phases, whereas BoP bursts dominated routine phases. These results demonstrate that frontal beta activity comprises at least two rhythmically distinct regimes linked to different levels of cognitive control. Our dual mode framework refines mechanistic models of transient oscillations and underscores the significance of burst waveform diversity for flexible brain function.

## Introduction

Rhythmic processes measured with magneto/electroencephalography (M/EEG) and local field potentials (LFPs) critically regulate neural network dynamics and cognitive functions (Miller et al., 2018). Capturing these rhythms at sub-second timescales reveals a burst-like organization: beta- and gamma-band signals often manifest as brief, discrete events rather than continuous waves (Feingold et al., 2015; Lundqvist et al., 2016; Sherman et al., 2016; van Ede et al., 2018; Little et al., 2019). Framing oscillatory activity as transient “bursts” thus provides a mechanistic lens for understanding neural processing, enabling precise characterization of burst timing, waveform diversity, and functional roles (Shin et al., 2017; Quinn et al., 2019).

Beta-band bursts in the 15–29 Hz range exhibit striking heterogeneity, from sustained multi-cycle oscillations lasting more than three cycles (Lundqvist et al., 2016) to ultra-brief transients under a single cycle (Sherman et al., 2016; Shin et al., 2017). This waveform diversity, highlighted by Szul et al. (2023), implies that beta bursts arise from distinct network mechanisms rather than a single stereotyped process. Precisely mapping the full spectrum of beta-burst waveforms will deepen our understanding of how these transient events contribute to neural computations within the same recording and frequency band. Importantly, it cannot be assumed that all of the burst-like phenomena in the beta band have the ***same*** role with respect to cognition, and this is what we seek to test in the current work.

Burst detection has traditionally relied on a spectral power threshold to identify high-amplitude transients—bursts of power (BoP). While BoP effectively captures strong events, it does not account for rhythmic stability or the internal structure of lower-power but rhythmically consistent bursts. Moreover, power-threshold methods can yield similar spectra from markedly different underlying signals (Jones, 2016), indicating that amplitude alone is insufficient to characterize transient beta activity. Recent improvements—adaptive thresholding and noise correction—have enhanced the specificity of BoP detection (Szul et al., 2023). Complementary techniques such as lagged coherence have also been used to quantify oscillatory stability, revealing additional structure in transient beta bursts across cognitive and sensorimotor contexts (Fransen et al., 2015; Little et al., 2019; Rayson et al., 2023; Szul et al., 2023).

To capture rhythmic structure directly, we introduce bursts of consistency (BoC), which detect events only when they exhibit stable oscillatory structure over multiple cycles. Adapted from the cycle-by-cycle framework (Cole and Voytek, 2019), BoC enforces thresholds on amplitude consistency, period consistency, cycle monotonicity, without reliance on relative power. We tuned these parameters through simulations in realistic noise to ensure reliable detection of genuine oscillatory bursts while minimising spurious events.

Importantly, BoP and BoC detections can overlap: some events satisfy both high-power and multi-cycle criteria, indicating they possess both high power and sustained rhythmicity. This overlap shows that each method captures distinct facets of beta activity—BoP uncovers brief, high-amplitude transients, while BoC isolates sustained, rhythmically regular bursts. Combining both approaches thus provides a more comprehensive characterization of beta-burst subtypes, and this characterization may help us to better understand the functional relevance of beta bursts, and in particular to reveal any diversity in functional roles..

Applying both methods to existing recordings from mouse barrel-cortex LFP (Shin et al., 2017) and macaque lateral-prefrontal LFP and ECoG (Quilodran et al., 2008; Stoll et al., 2016), we identified two reproducible beta-burst types that are largely independent. BoP bursts have a large amplitude peak and little apparent rhythmicity around that peak, whilst BoC bursts are more classically rhythmic or sinusoidal over several cycles. In the macaque data we show that in a well-established task in which animals are consistently cycling between phases of high (search) and low(repeat) cognitive-control demands every few trials, BoC bursts were more frequent during demanding search phases, while BoP bursts predominated during repetition phases. This double dissociation reveals that frontal beta dynamics comprise at least two rhythmically distinct transient regimes, each tied to different functional states.

## Results

We established two contrasting methods to detect bursts of activity in neural signals. First a refined BoP approach building on the widely reported six factors of median (FOM) method (Feingold et al., 2015; Shin et al., 2017), but restricting it to a 2-cycle minimum to cut down on possible spurious detections discovered in preliminary simulations (S1 Fig, also see Langford et al., 2025 for a discussion on the pitfalls of BoP method). which we refined in preliminary simulations (S1 Fig). Second a BoC method derived from the cycle-by-cycle approach (Cole and Voytek, 2019) with detection parameters selected for optimal burst non-detection rate in 1/f noise. With each method we extracted the timing of the bursts, and then associated that burst with frequency and amplitude properties using the same instantaneous frequency and amplitude estimates between 15 and 29 Hz for each method – thereby ensuring reliable comparison of these features between methods. If a burst occupied half the same temporal space in both methods, we classified it as a BoP/C burst common to both methods, whilst other bursts were described as unique to one class or the other.

### Cycle consistency reveals unique burst characteristics

We applied these approaches to mouse data (Shin et al., 2017) as a first methodological test. Looking at burst amplitude, duration and frequency, we observed that the BoP and BoC methods led to different outcomes for burst detection (Fig 1). The BoC method detected bursts with slightly lower frequencies (M=20.2 Hz, SD=0.457) to those detected by the BoP method (M=20.9 Hz, SD=0.28); t(9) = -5.69, p < 0.001. The durations were also shorter for the BoC (M=0.12 s, SD=0.009) compared to the BoP (M=.146 s, SD=0.007); t(9) = -7.98, p < 0.001. Finally, the amplitudes were lower for the BoC (M=0.54 normalize mV, SD=0.06) compared to the BoP (M=.631 normalize mV, SD=0.014); t(9) = -4.19, p < 0.01. Amplitude differences were, of course, expected given the threshold cutoff in the BoP method, and its absence in the BoC. While the BoP method detected a large number of bursts (810), the BoC method detected fewer bursts (134). Strikingly there was little intersection (48) between the two methods (BoP/C), suggesting that we are detecting independent phenomena with the two approaches. When we looked at individual burst waveforms, it appeared that the BoCs have more consistent peaks and troughs throughout the duration of the bursts, whereas the BoPs tend to be dominated by a singular large peak, with very little consistency over the duration (Fig 2, with raw data between burst onset and offset min-max normalized to focus on rhythmic differences). The BoP/C bursts tend to, as expected, follow the BoC bursts in their consistency, and we know that they are also above 6 FOM threshold, and so have high amplitudes.

**Fig 1.**
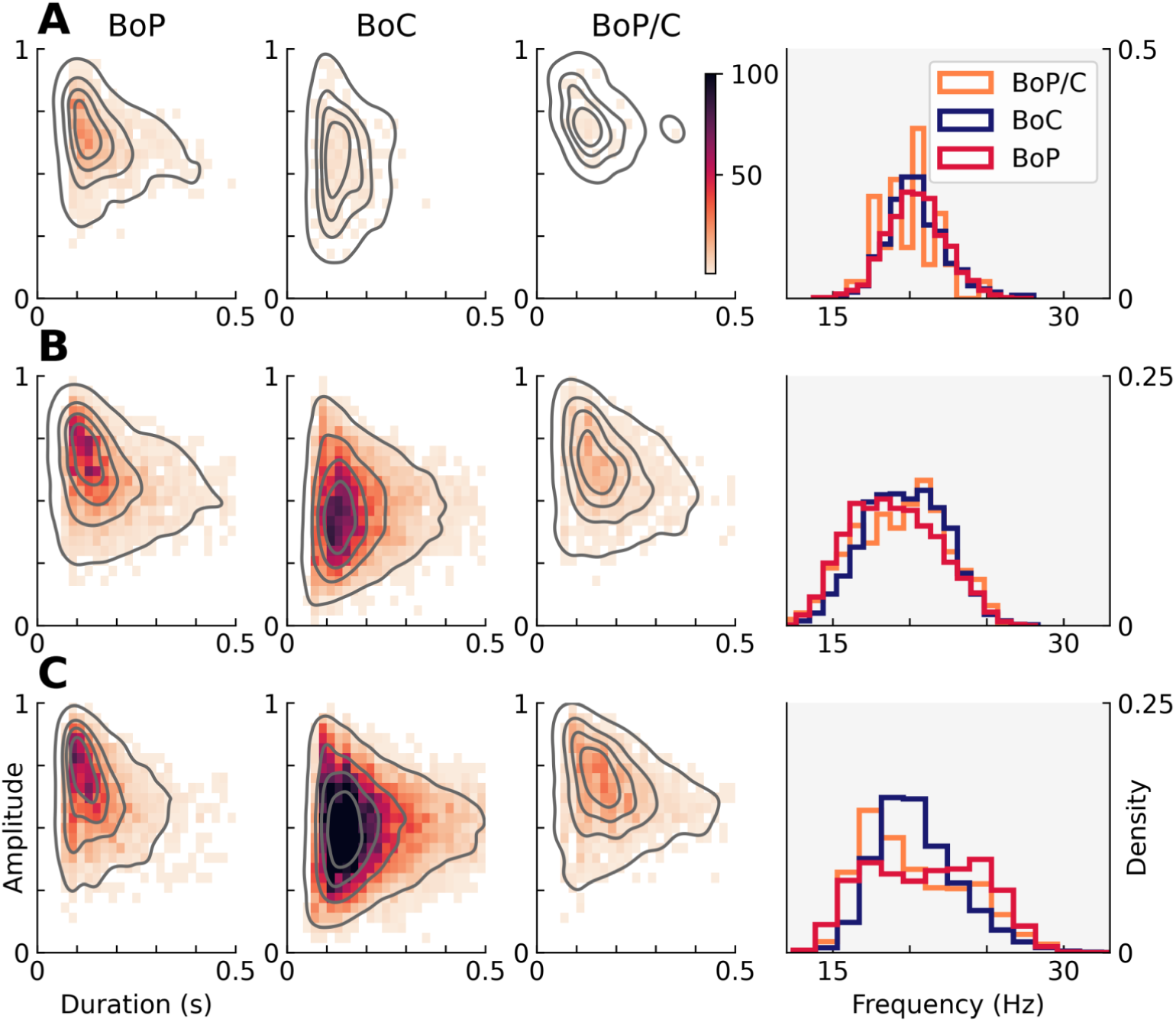
Scatter plots of the duration and amplitude for the BoP, BoC, and BoP/C bursts detected in the three leftmost panels for (A) mouse LFP, (B) macaque LFP, and (C) macaque ECoG. Frequency histogram of each method in the rightmost panel.

**Fig 2.**
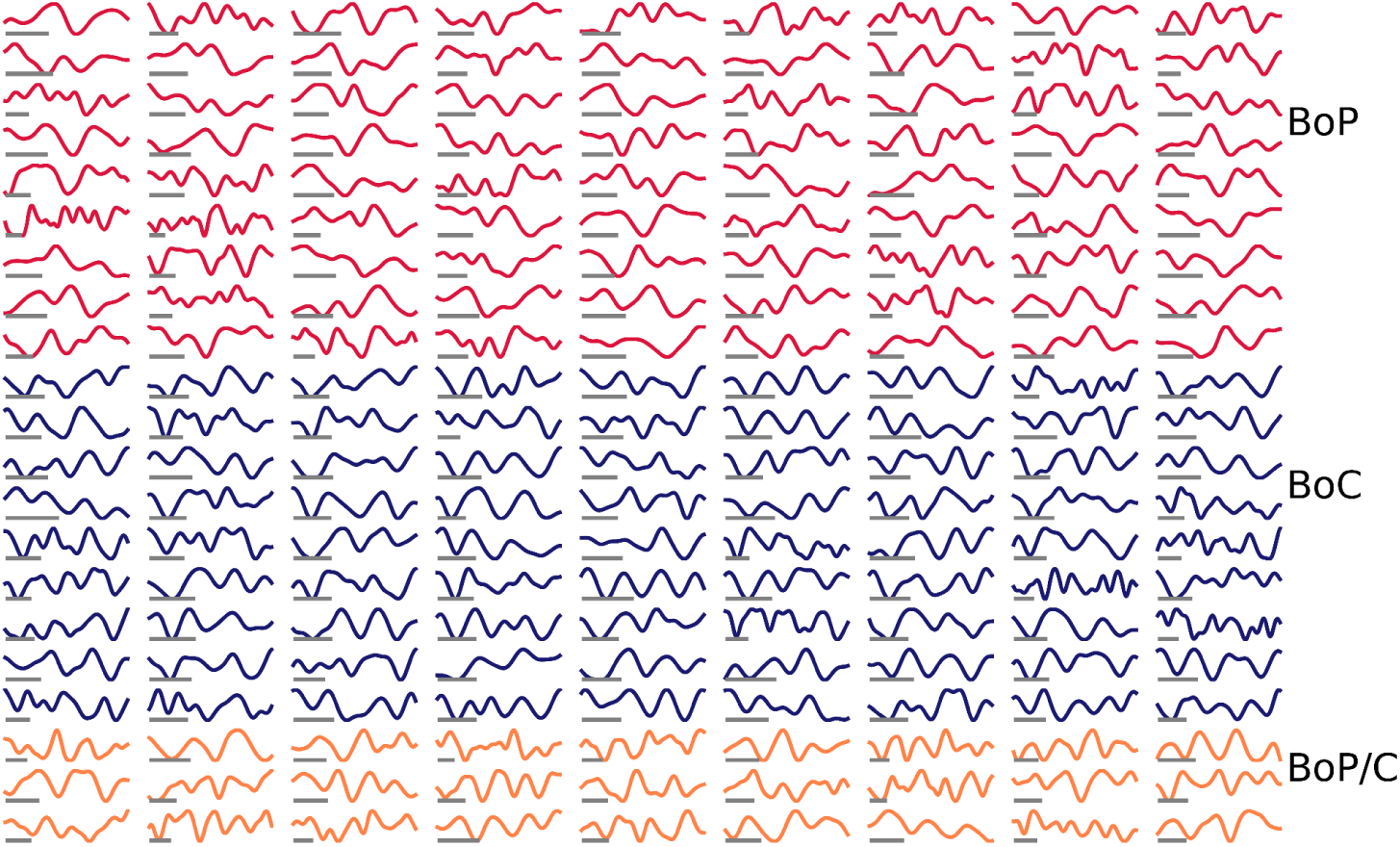
Random selection of (<7 cycle) BoP bursts (top), BoC bursts (middle), and BoP/C (bottom) for the mouse LFP data. Grey bar at bottom left of the graphic is 1 cycle at that particular burst’s median frequency (calculated using the relevant instantaneous frequency time course).

We performed equivalent analyses on macaque LFP data from the lateral prefrontal cortex (PFC) recorded during the delay period of trials of a search-repeat task (Figure 1B), using data previously acquired by the lab in two monkeys (Quilodran et al., 2008). This task required the monkeys to *search* an array of 4 targets using trial and error to find the rewarded target, and then *repeat* the rewarded choice for several trials. The delay was the period of time at the start of the trial prior to the selection of an option.

When looking at the properties of the resulting bursts, duration was not different between the BoP and BoC methods. Unsurprisingly, the amplitude was lower in the BoC (Monkey M BoC: M=0.45 normalized mV, SD=0.02 and BoP: M=0.62 normalized mV, SD=0.02; t(14) = -23.14, p < 0.001. Monkey P BoC amplitude: M=0.48 normalized mV, SD=0.03) and BoP: M=0.64 normalized mV, SD=0.05 ; t(13) = -8.25, p < 0.001). The frequency was higher in the BoC (M=18.8 Hz, SD=0.53) than in the BoP (M=18.4 Hz, SD=0.42; t(9) = 4.68, p < 0.001). In contrast to the mouse data, the BoC count for both monkeys (monkey M = 3139; monkey P = 2660) was higher than the BoP count (monkey M = 1795; monkey P = 1005). Yet like the mouse data, the shared BoP/C burst count was significantly lower than the unique burst counts, although the difference was proportionally smaller (monkey M = 511 monkey P = 403), again supporting the notion that we are detecting independent phenomena.

Finally we applied the methods to ECoG data (Stoll et al., 2016) from the lateral PFC of one monkey recorded over multiple instances of a similar task (Figure 1C). The durations were longer for the BoC (M=0.19 s, SD=0.006) compared to the BoP (M=.144 s, SD=0.007); t(24) = 21.10, p < 0.001. As observed in the other datasets the amplitudes were lower for the BoC (M=0.50 normalize mV, SD=0.01) compared to the BoP (M=.65 normalize mV, SD=0.02); t(24) = -33.65, p < 0.001. The frequency is lower in the BoC (M=20.5 Hz, SD=0.18) than in the BoP (M=21.0 Hz, SD=0.34); t(24) = -6.35, p < 0.001. Similar to the macaque LFP, but dissimilar to the mouse, there were many more BoC (12,443) than BoP (3112), and also notably fewer BoP/C (1633).

In conclusion, whilst there was overlap between the two methods in the monkey data, the level of bursts unique to one detection method was striking. The fact that the two methods provide for the most part non-overlapping events as bursts suggests that the BoP and BoC methods are extracting different neurophysiological events. Given the differences in detection principles some differences were to be expected, but the extent of the separation between the two is striking.

### Evaluating burst type differences as a continuum

Given the scarcity of BoC bursts in the mouse data and the relatively small overlap between BoP and BoC bursts across all datasets, we investigated whether this limited intersection was purely a consequence of parameter selection. To systematically assess this, we tested 512 parameter sets in the initial session of the mouse dataset, spanning 208 trials that originally contained 86 BoP bursts. The parameters—amplitude consistency, period consistency, and cycle monotonicity—were systematically varied from 0.01 to 0.7 in steps of 0.1, generating a wide range of detection criteria. To quantify the overall strictness of each parameter set, we introduced a severity metric, defined as the sum of the three parameters. For each set, we calculated the number of BoC bursts that temporally coincided with any of the 86 BoP bursts, yielding a percentage overlap per parameter set, i.e., the percent of BoP/C at each severity.

Visualizing these results in Fig 3 allowed us to evaluate the level of leniency required to achieve significant overlap between BoP and BoC bursts, relative to the leniency of the original parameters. Our findings indicate that even under highly permissive parameter settings, and a permissive overlap threshold, overlap between the two burst types remained constrained. At the most lenient setting (all parameters at 0.01), the intersection reached only around 50%. Notably, at these extreme parameter values, the BoC method identified a substantial number of bursts (∼1.5 per trial), not only in the mouse data but also in simulated noise, suggesting that it was detecting events with weak or ambiguous oscillatory characteristics. This implies that the limited overlap observed at less severe settings likely does not reflect a true physiological relationship between BoP and BoC bursts but rather overly permissive parameter settings, and thus there is a real division between BoP and BoC bursts., i.e., these findings support the existence of distinct burst types rather than reflecting a shared oscillatory phenomenon. Even in the case that we were more permissive, there still wouldn’t be a substantial BoP/C count.

**Fig 3.**
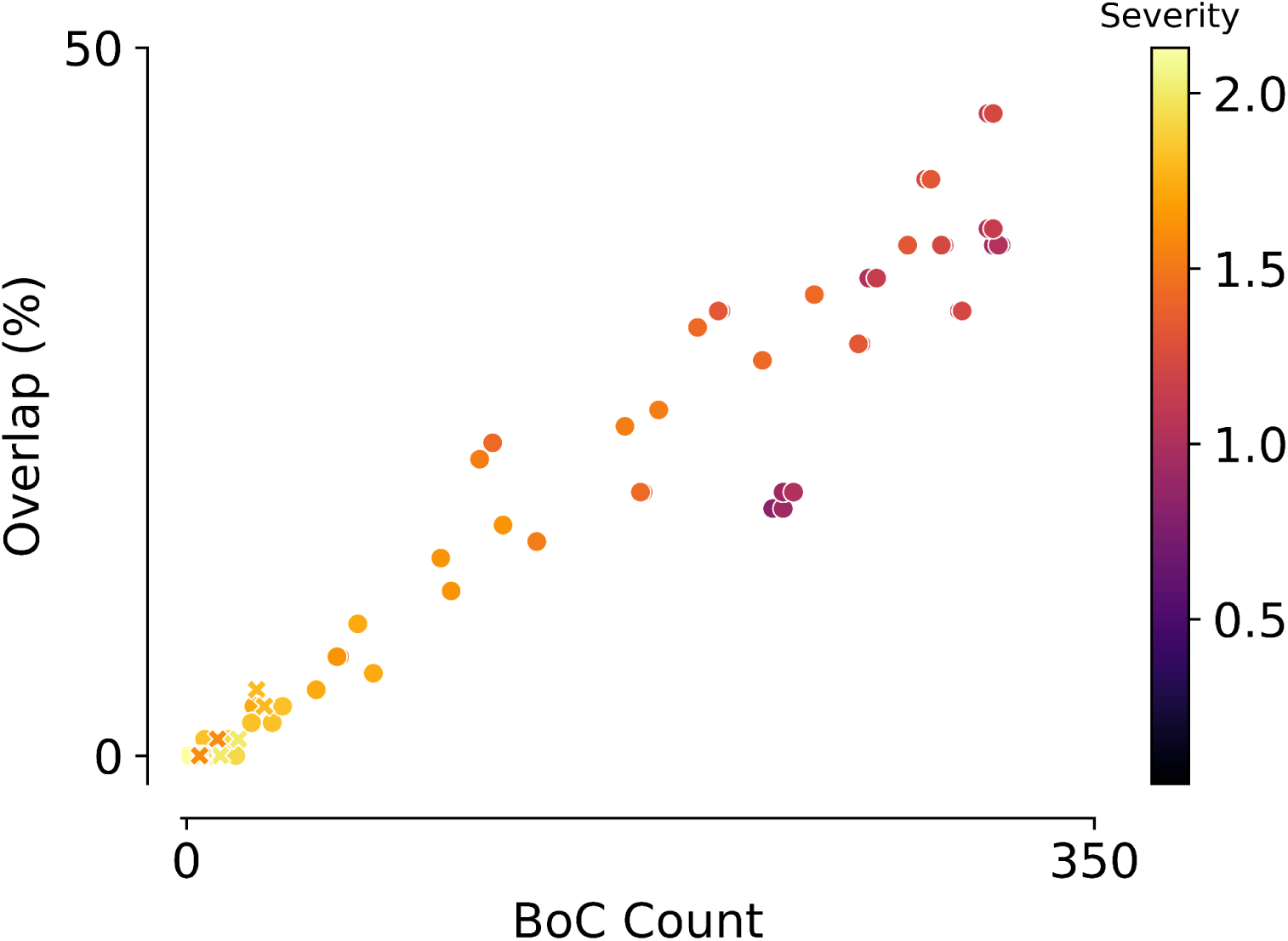
Percentage of bursts recovered by both methods over total BoC count and parameter severity for the initial session (208 trials) of the mouse data. The crosses are to highlight the specific set of parameters used in the study presented here. Severity is the sum of the BoC method parameters for a given parameter set.

### Bursts of consistency show a more visually rhythmic signature

Here, we sought to provide a qualitative description of the features of the two emerging classes of burst revealed by our two analysis approaches. Notably, could we show a general effect of the rhythmic differences we noticed in the individual bursts of the mouse LFP. To achieve this, we created averaged burst-related potentials for each dataset and method. We studied a subset of bursts in the middle of the frequency range considered above, and aligned each burst at the greatest absolute amplitude within the burst, plotting the resulting averaged waveform as a burst-related potential in Fig 4.

**Fig 4.**
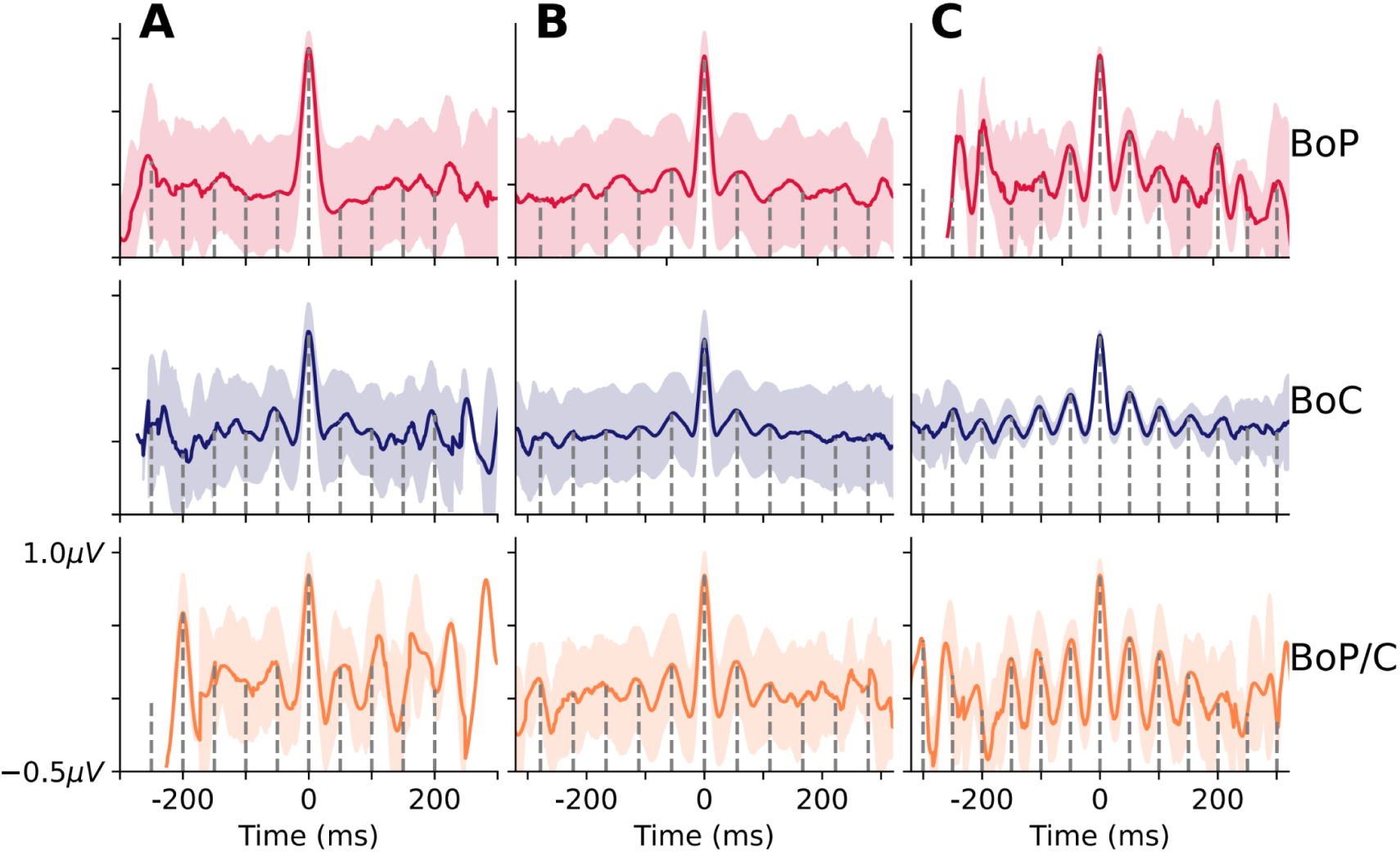
Burst Related Potentials. Raw data are aligned at the greatest absolute amplitude of each burst for mouse (A), LFP (B), and ECoG (C) across the BoP (top), BoC (middle), and BoP/C (bottom) methods. The gray dashed vertical lines denote the duration of 1 cycle at the median frequency aligned with the burst. Notably, even in the average, BoC exhibits more peaks aligned with the gray line—visually suggesting a more pronounced rhythmic structure than BoP—while BoP/C shows high peak alignment accompanied by greater amplitudes.

In both sets of monkey data bursts derived from the BoC method are particularly rhythmic, with peaks occurring at the median frequency time point (gray dashed lines) over multiple cycles before and after the peak. In contrast, bursts derived from the BoP method remain rhythmic at this frequency for notably fewer cycles, whilst the absolute value of the peak is larger – confirming observations from the individual burst plots in Fig 2. It should be noted that in this analysis we have selected the highest absolute amplitude for alignment, so by definition the waveform has to decrease from the center. But only if it is rhythmic will it then again increase and have a local peak at ∼1 cycle. These analyses suggest therefore that the form of the bursts detected by the two methods differs significantly - BoPs were more likely to be transients with a large peak and little rhythmicity in cycles around that peak; BoCs were more rhythmic and sinusoidal over several cycles.

### Functional specificities of different burst types

The two separate classes of burst in the beta band described above appear to represent different neurophysiological phenomena – they occur independent in time to a large extent, and they have notably different rhythmic properties. As such we may be observing two independent mechanisms within the same frequency band. A key question is therefore whether these two types of burst have different *functional* roles. We were able to test this in the macaque data because the LFP and ECoG were acquired while animals were regularly shifting between different behavioral phases, searching by trial and error and repeating rewarded responses (Quilodran et al., 2008; Stoll et al., 2016). These previous studies form part of a body of work that shows how frontal neural dynamics prospectively control the switching every few trials (2-4) between search (S) and repetition (R) phases. Importantly, trials in S and R phases are identical - what differentiates them is the monkey’s knowledge about what they must do in the upcoming trial. In the case of the beta oscillations studied here, a significant decrease in trial averaged beta power in the delay period is observed when the animals move from search to repeat trials (Stoll et al., 2016; Wilson et al., 2016). This shift is also reflected in single neuron firing in the cingulate cortex (Quilodran et al., 2008), gamma oscillatory power in cingulate and lateral prefrontal (Rothé et al., 2011), and attractor dynamics of the frontal population (Enel et al., 2016). As such we asked whether the burst classes described above both showed this property of shift between search and repetition, and how.

For each trial we calculated the number of bursts derived from BoP and BoC detection methods and for each behavioral phase in both the LFP and ECoG. Fig 5 illustrates the commonly reported burst rate (average per trial) of each burst type. In addition to a difference in overall rate between BoP and BoC, this clearly highlights differences between S and R phases, in line with expectations from prior studies on averaged beta power (Stoll et al., 2016). Notably, the striking observation is that BoP and BoC show opposite patterns during these phases, suggesting potential divergent functional roles. To confirm this holds across each animal, even though prevalence of S and R trials varies somewhat between sessions, we plotted the ratios of S/R bursts for the two burst types against the ratio of S/R trials found in each recording unit. Fig 6A shows a single point for each recording unit (i.e., location, depth, monkey) in the LFP data, whilst Fig 6b shows a point for each recording session of the ECoG data. If the bursts occur equally across S and R trials in the neural signal we would expect the S/R burst proportion to align closely with the S/R proportion found in the trial structure, represented by the diagonal line. Moreover, if BoP and BoC are encoding the same behavioral information we would expect the pattern to be the same for the two burst types. Fig 6 brings out the striking difference between the two classes of burst – BoC bursts are more likely to be found in the search phase, and BoP bursts more likely in the repetition phase. We specifically tested this by using a binomial test with the null hypothesis of probability=0.5, effectively testing if the number of S/R bursts above or below S/R trials occur with equal frequency. In the case of BoP bursts we found that for both monkeys S/R was significantly different than the null (p_M_ < 0.001 with 13 of 15 S/R bursts below S/R trial, and p_p_ = 0.028 with 11 of 14 S/R bursts below S/R trial), and this result was further strengthened with the ECoG data (p_ECoG_ = 0.022, with 18 of 25 S/R bursts below S/R trial). For the BoC we observed a nearly completely opposite effect - with the S/R burst falling above the S/R trial. While the test for one of the LFP monkeys was not significant at p=0.05 (p_M_ < 0.059, 11 of 15 S/R bursts above S/R trial), the other animals combined strengthened this observation; p_p_ = 0.028, 11 of 14 S/R bursts above S/R trial, and p_ECoG_ < 0.001, with 22 of 25 S/R bursts above S/R trial. This means that BoC bursts are more likely to be found in the search phase, and BoP bursts more likely in the repetition phase.

**Fig 5.**
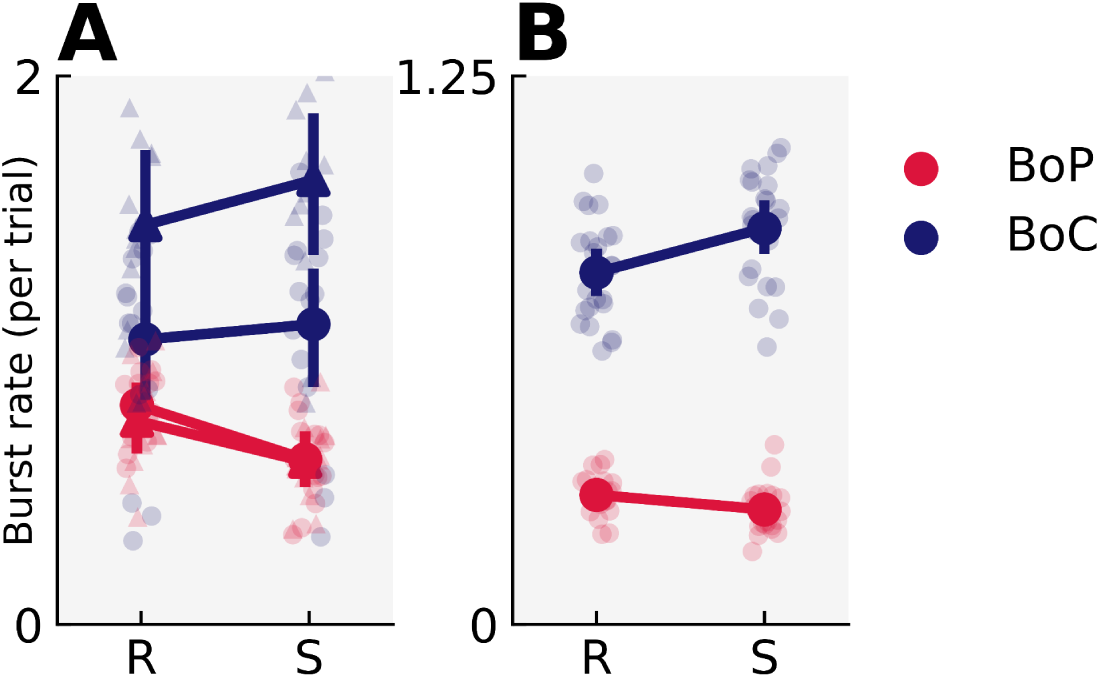
Burst rate of BoP and BoC in the search (S) and repetition (R) phases of the problem solving task for LFP (A) and ECoG (B) data. LFP data is shown for both monkeys, with monkey Marco shown as a triangle, and Pablo as a circle.

**Fig 6.**
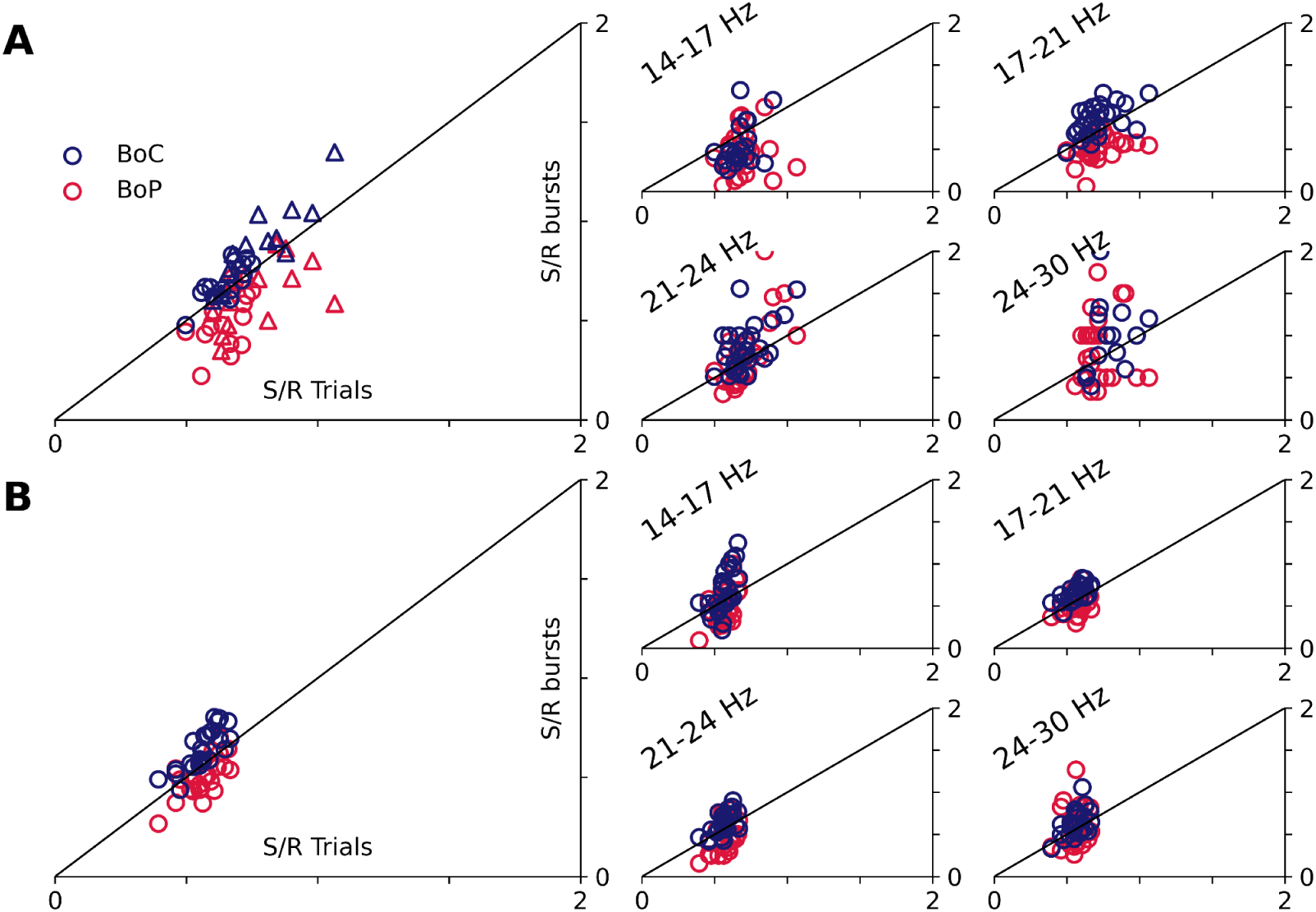
Distribution of BoP and BoC in the search and repetition phases of the problem solving task. The ratio of search and repetition trials per LFP (A) recording by ratio of search and repetition bursts found in that unit from 15-29 Hz and split between four different frequency ranges. In the leftmost plot the data is shown split by monkey, with Marco being represented by the triangle and Pablo by the circle. (B) is the same procedure as (A) but with the ECoG data, and with the recording unit being a session.

In addition, we verified that differences in the frequencies of the burst types were not accounting for the difference with behavior or related to frequency heterogeneities (for example the double frequency peak observed in the right panel of Fig 1). The right-hand column of Figure 6 shows that this was not the case. For the LFP data both the BoP and BoC ratios seem to be driven by bursts happening between 17 and 21 Hz for both monkeys (p<0.05 for BoC and BoP), 14-17 Hz for BoP in one monkey (p_M_ < 0.05), and in the same monkey 21-24 Hz for BoC (p_M_ < 0.05). In the ECoG it seems to be driven by the two middle frequency bins (17-21 Hz, p_EcoG_ < 0.05; 21-24 Hz, p=0.053), and not the lower, or higher. Importantly, therefore, the behavioral effect in bursts from either method was not purely driven by bursts in other frequency bands.

So the different forms of burst detected with the two methods have different relations to behavioral phases. Burst analysis provides an analytic resolution beyond the average across trials, and so we can also study individual burst properties and their variation between search and repetition. Fig 7 therefore takes the 3 major individual burst properties of amplitude, duration, and frequency, to test whether we see search-repetition differences here in addition to the differences in distribution described above, using paired samples t-tests within each method. The figure also brings out the initial BoP vs BoC property differences already tested in Fig 1.

**Fig 7.**
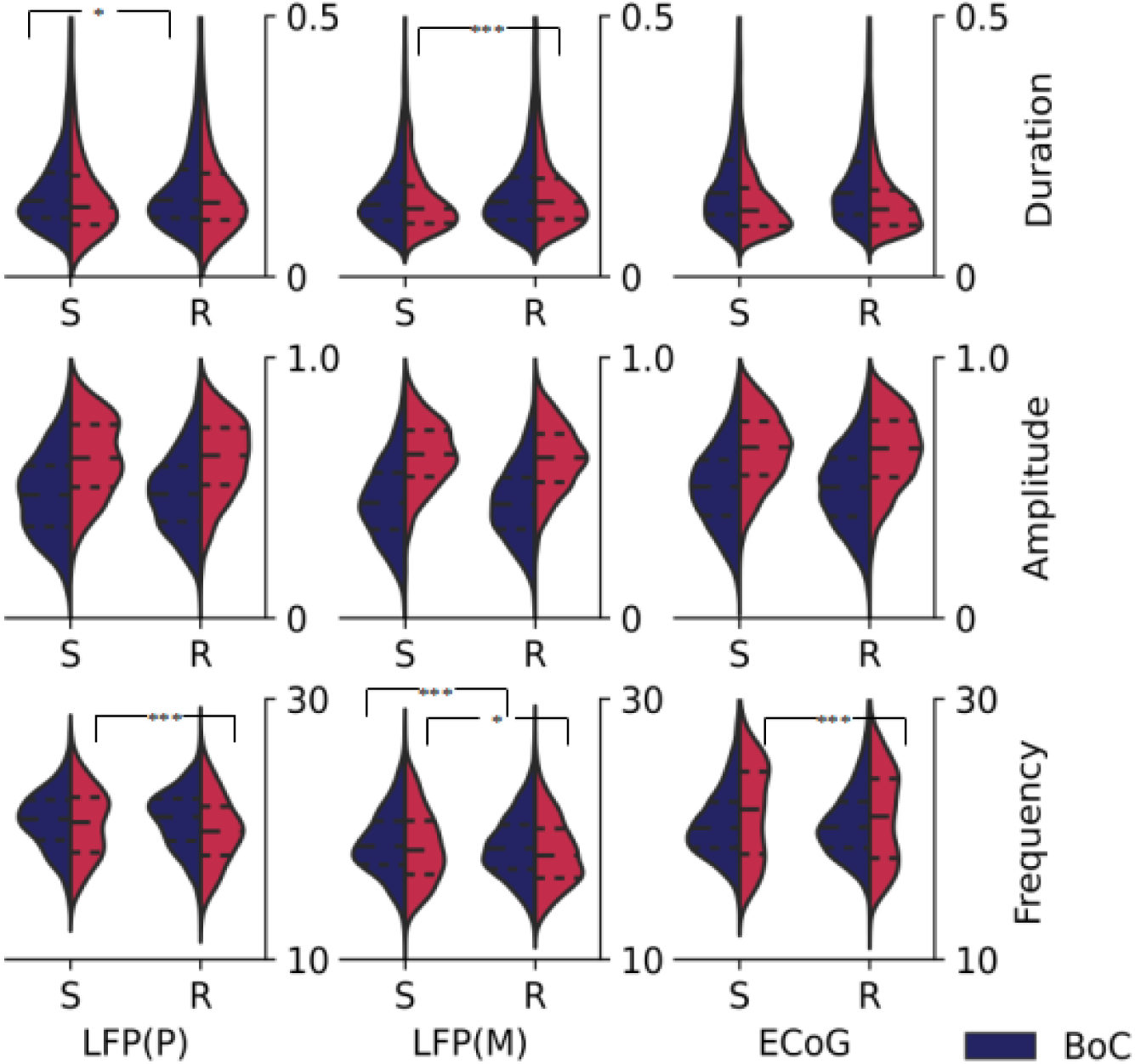
Differences in the burst characteristics in the search and repetition phases of the LFP and EcoG split by BoC and BoP bursts. *** = p <0.001, * = p<0.05

We found no differences in amplitude between search and repetition, for either BoC or BoP in any of the datasets (Fig 7 middle line), despite a clear difference in average power previously described (Stoll et al., 2016). This *suggests* that the average power differences are ***not solely*** related to burst amplitude differences – potentially a surprising result in that an intuitive interpretation of increases in *average* power in a frequency band is that amplitude of components at those frequencies has increased. We suggest that the average changes in power can be accounted for by changes not only in burst power, but in burst count, and burst duration. This contrast is one that will be important to develop in future analyses seeking to understand the contribution of such mechanisms to average power.

We also observe a consistent frequency shift between search and repetition for BoP only, but only inconsistent variations in burst duration. Specifically For monkey P we found differences in the BoC durations, being less in S (M=0.17 s, SD=0.008) than R (M=.17 s, SD=0.012); t(13) = -2.49, p = 0.03. BoP frequencies were higher in S (M=20.8 Hz, SD=1.29) than R (M=20.2 Hz, SD=0.012); t(13) = 3.21, p<0.001. For monkey M we found differences in the BoP durations, being less in S (M=0.15 s, SD=0.014) than R (M=.17 s, SD=0.014); t(14) = -3.19, p <0.00. As for monkey P, monkey M’s BoP frequencies were higher in S (M=18.7 Hz, SD=0.82) than R (M=18.3 Hz, SD=0.33); t(14) = 2.27, p=0.04, but there was also an effect in the BoC frequencies with S (M=19.0 Hz, SD=0.52) compared to R (M=18.7 Hz, SD=0.60); t(14) = 2.97, p=0.01. For the EcoG only the BoP frequencies were higher in S (M=21.3 Hz, SD=0.52) than R (M=20.9 Hz, SD=0.39); t(24) = 3.66, p=0.001.

So, the two types of burst are differently distributed between search and repetition, but they also show different modifications of their properties between the behavioral phases – the different neural dynamics reflect the engagement of neural mechanisms specific to each behavioral phase. Such analysis demonstrates how future studies can begin to build up a detailed picture of the differing mechanisms, moving towards a full description of cognitive state-specific neural dynamics.

## Discussion

Single trial analysis detailing the transient nature of beta oscillations in the frontal cortex has become an important approach to study how oscillatory signals can be interpreted and understood mechanistically within neural networks. The form of the bursts described in the literature varies significantly, and so here we apply two complementary yet contrasting analyses and burst extraction approaches to provide evidence that within the frontal beta band there are at least two differing transient phenomena in data across species and recording modality. Using two different criteria for burst definition should inevitably generate differences in detected properties, but the extent of these differences is striking - the level of overlap in the phenomena detected is surprisingly low. This leads us to propose that there are in fact two different phenomena occurring in the beta frequencies in the recordings that we have studied across species and recording modalities - the short transients (BoP) which resemble the phenomena previously described by several groups (e.g. Sherman et al., 2016; Shin et al., 2017), and the more oscillatory signals lasting several cycles (BoC) but with lower peak power. BoP and BoC show opposing distributions in two behavioral phases of a search-repetition cognitive task, showing that the two phenomena have differing functional as well as physiological properties.

One might speculate that these two phenomena, with different relationships to behavioral measures in our study, have different physiological generators. For example Sherman et al (2016) have described a model where the transient beta events are driven by a short distal input that arrives, presumably, from outside the cortex. In contrast, the longer, more sinusoidal and consistent bursts resemble more the phenomena derived from a local circuitry with oscillatory events emerging from interactions and conduction delays between pyramidal cells and interneurons (e.g. Lundqvist et al., 2011).

Given their different spectral properties, it is not necessarily surprising to find more than one phenomenon of transient event at roughly the same beta frequency- one can imagine that the network detects and responds to lower power sinusoids and high power transients very differently, and so their occurrence at very similar frequencies is unlikely to lead to confusion. The next stage of this project is to understand this process and thereby use the data here to dig further into local mechanisms. The results here are therefore a significant advance in our quest to understand how the cortex may be using such signals.

The results are also significant from a methodological standpoint as well. Our aim here was not to provide an exhaustive comparison of burst detection methods, indeed other authors have begun to provide this in several contexts (Cho & Choi 2023; Ardelean et al., 2023). But our focus on comparing bursts determined by two different features lays bare the importance of the method by which transient events in local field signals are detected – indeed in our hands the behavioral result (and therefore inferred behavioral role of the beta bursts) is ***reversed*** between the two detection methods. In addition, we have demonstrated that changes in trial average power do not necessarily result from changes in single trial or burst amplitudes, as has often been inferred (Fig 6). These two results demonstrate the power of the burst analysis approach to progress our understanding of these phenomena in neural signal. Future work will seek to capture the two phenomena with a signal analytic approach, and we recommend going forward that both power and consistency criteria are considered in detecting local field bursts.

Yet, burst amplitude, duration, rate, and average power are interrelated, and our analysis does not explicitly model how burst amplitude may contribute to these effects over time. Prior work in sensorimotor beta suggests that post-movement beta rebounds involve both burst amplitude and rate changes, but the present task differs in cognitive demands and does not directly involve motor execution. Given this, while our findings suggest that burst amplitude *alone* does not fully account for the task-phase average power differences, we acknowledge that further work is needed to fully disentangle the relative contributions of burst rate, duration, amplitude, and overall rhythmic activity across task phases.

The observed dissociation between BoP and BoC bursts in our study aligns with the distinct cognitive demands of search and repetition trials. In search trials, animals must actively explore possible solutions, while in repetition trials, they must simply repeat the response to the rewarded location that has been determined by the search. This difference in cognitive load and strategic engagement is reflected in the contrasting prevalence of BoC (search-dominant) and BoP (repetition-dominant) bursts. Notably, this pattern mirrors prior findings that frontal beta oscillations are modulated by cognitive control and decision uncertainty (Stoll et al., 2016; Wilson et al., 2016), but also more widely it follows findings about changes in spiking activity and gamma oscillations in prefrontal and cingulate cortex that reflect this state shift between search and repetition phases (Procyk and Goldman-Rakic, 2006; Quilodran et al., 2008). Our approach contrasts with studies of sensorimotor beta bursts, which often examine trial-to-trial burst-behavior correlations in tasks involving motor execution (e.g., Szul et al., 2023). In such tasks, beta bursts are typically linked to movement preparation, execution, or inhibition, making it reasonable to assess bursts at the single-trial level. However, in a higher-order cognitive task such as ours, search and repetition represent distinct cognitive states rather than isolated motor events. Analyzing how burst types are distributed across these task phases provides insight into state-dependent neural dynamics. The contrast between BoC and BoP bursts suggests that beta activity in the frontal cortex may serve different computational roles depending on task demands, potentially reflecting distinct network states for **exploratory decision-making.** Future work could explore whether similar burst differentiation exists in other cortical regions, including the sensorimotor system.

The contrast between the two methods can be questioned on the basis of chosen parameters - it would presumably be possible to parameterize the BoC method to be less conservative and capture some elements of the BoP bursts described here. But the strong result here is that the BoC and BoP bursts occupy different space altogether in the behavioral paradigm, which seems to suggest they are functionally distinct phenomena as captured by the analyses presented here. It is also worth noting that both methods are capable of picking up - at least - two cycle bursts, as we used this as the lower bound. The BoP method is actually capable of detecting slightly shorter bursts because of the smoothing involved in TF decomposition, it will also tend to report a burst as being slightly longer for this same reason. In this sense, it is not possible to make a true direct comparison of burst length related to the raw signal, and modifications of burst length should be considered within-method only; and should be the subject of further study.

The contrast between BoC and BoP bursts in the behavioral paradigm are expressed across the beta band, and in very similar sub-bands within the classical beta frequencies (Fig 5, right), despite there being systematic differences between overall distribution of frequency for the two methods (Fig 1, right). Again, there is an element of frequency smoothing to the BoP method that makes direct frequency comparisons complicated without re-projecting those bursts onto raw signals as used in the BoC approach. Understanding the role of frequency of individual events in the beta band, in order to be able to interpret the meaning of averaged frequency, should be a strength of the burst analysis approach, and a detailed analysis of burst event frequency will be the subject of future work.

### Regional and Spatial Considerations in Beta Burst Dynamics

While our study focuses on frontal beta bursts, recent work suggests that sensorimotor beta bursts may differ in structure and function. For example, sensorimotor beta bursts remain transient across conditions, even when sustained motor output is required (Echeverria-Altuna et al., 2021). In contrast, our frontal recordings reveal BoC bursts with more rhythmic properties, potentially resembling more classical oscillatory activity. This suggests that frontal circuits may support multiple burst regimes, a distinction that may not necessarily generalize across brain regions.

Further highlighting potential regional differences, the rodent data—recorded from somatosensory cortex—showed relatively few BoC bursts compared to the primate frontal recordings. This scarcity may reflect differences in burst organization across cortical areas, though species- and task-related factors must also be considered. More broadly, understanding how beta burst dynamics vary across brain regions and cognitive contexts is a critical question in this field, to be addressed in the future through concurrent recordings in multiple regions and across multiple forms of task.

### Implications and Future Directions

Our findings demonstrate that beta-band activity in the frontal cortex is not a singular phenomenon but instead comprises at least two distinct burst types with opposing behavioral relevance. By distinguishing between high-power transients (BoP) and rhythmically stable bursts (BoC), we reveal an underlying structure in beta dynamics that would be obscured by traditional trial-averaged approaches. The fact that these burst types are differentially modulated by task phase suggests that they reflect separate functional mechanisms rather than arbitrary divisions within a continuum of activity. Moving forward, understanding how these burst dynamics emerge from neural circuits, how they interact with other oscillatory processes, and how they generalize across brain regions will be crucial. If beta bursts shape cognition and behavior in a structured yet transient manner, then decoding their precise temporal signatures may offer powerful insights into neural computation, cognitive control, and even clinical interventions targeting disrupted beta activity.

## Methods

The methods presented here are described in an IPython Notebook on github at https://github.com/langfordzd/burst_regimes/blob/main/walkthrough.ipynb and the data for the LFP and ECoG analyses can be retrieved at https://osf.io/qz59n.

### Burst detection methodology similar across data types

#### Bursts of power method

The BoP approach used here builds on the widely reported six factors of median (FOM) method (Shin et al., 2017), which we refined in preliminary simulations (Fig S1). We defined BoPs as contiguous surfaces above the 6 FOM threshold lasting at least 2 cycles at the median frequency.

This method utilizes a time-frequency analysis approach, where the raw signals are convolved with a complex Morlet wavelet at 7 cycles to generate a time-frequency map in the frequency range of 15 to 29 Hz. The power estimates at each frequency are then normalized across frequencies by dividing them by the median frequency at that specific frequency. To align with the current method used in the field, the method defines high amplitude events as any part of the time-frequency map that is at or above 6 times the median frequency. To address the issue of false bursts, the method further refines the definition by considering only bursts with temporal lengths that are over 2 cycles at the burst frequency, as this aligns with the lower duration bound of the BoC-method and effectively limits the number of false bursts.

#### Bursts of consistency method

The BoC method takes advantage of a recently developed approach known as the cycle-by-cycle method (Cole and Voytek, 2019). This framework was designed to characterize the presence or absence of a consistent oscillation in a time-domain signal. This determination is based on the definition of a periodic process, in which the voltage at time t+tau should be predictable from the voltage at time t, where the process has a period tau. In order for the future voltage to be reliably predicted from the past voltage, the amplitude and period must be conserved from cycle to cycle. This is done by examining the symmetry between amplitude peaks and troughs within a cycle (amplitude consistency), and the symmetry between the rise and decay of a cycle (period consistency) There are two further parameters we were interested in; the monotonicity of the cycles, and the number of cycles. We would like to be clear and point out that this work does not use the built-in amplitude thresholding method of the cycle-by-cycle Python package to detect bursts. The point of difficulty in the BoC method is in defining thresholds for these parameters in a principled manner. Our main contribution to this method is in developing a technique to find reasonable sets of parameters to detect bursts of oscillation in a non-arbitrary manner. First we analyzed 128 possible parameter combinations (between amplitude-consistency, period-consistency, cycle-monotonicity, and cycle count), to identify the most suitable ones that yielded a reasonable burst non-detection rate in 1/f noise. This involved two different steps of regularization; the first being to only use amplitude-consistency, period-consistency, and cycle-monotonicity values from 0.2 to 0.8 (in steps of 0.2, 64 unique combinations) with the number of cycles to be 2 or 3 (bringing the total to 128 unique parameter combinations). The second step involved pruning this four dimensional grid to only include parameter combinations that would only rarely classify a noisy signal as an oscillatory burst. To do this we first simulated 1000 trials of 1 second coloured noise and calculated the burst count for each trial. We then selected 100 trials randomly, and with replacement, 1000 times and calculated the mean over these selections. We aimed to pick sets of parameters that would only produce, on average, one burst per ten trials in coloured noise. To further prune it we then excluded parameter sets that would only produce one burst every 30 trials, because these would be too conservative and in practice would not detect bursts (either in the simulation or the electrophysiology). To pick among these regularized sets of parameters we took the set with the highest trial-averaged correlation between the instantaneous amplitude and the temporal profile of the recovered bursts (binary coded; 0 was not bursting and 1 was bursting). Details of the parameters that provided this solution can be found in the iPython notebook accessible through Github describing the analyses presented here. The final goal of this work is to use this method to detect bursts of oscillation in a non-arbitrary manner, by using well-defined thresholds for the parameters, which are determined by simulation and analysis.

#### Burst property extraction

To further define the bursts in both methods we determined the median frequency and amplitude within the start and end timepoints for both methods using the instantaneous frequency and amplitude estimates between 15 and 29 Hz. This ensured that we had a relatively independent measure of frequency and amplitude for comparisons between methods, as the methods each only defined a burst by their timings (i.e., the burst frequency of the BoP method was not taken from the same TF decomposition). Importantly, if a burst occupied half the same temporal space in both methods, without doubling it, we classified it as a BoP/C burst common to both methods.

#### Burst related potentials

We accomplished the burst related potentials by first subsetting the bursts that fell within a smaller frequency window around the center of the frequency distribution, so while there is not much difference in the frequencies to look at temporal alignment, we capture a large portion of bursts. For the mouse we took this window to be 19-21 Hz, for the macaque LFP we used 16-20 Hz, and finally for the EcoG we chose 19-21 Hz. We then aligned each burst in each method at the greatest absolute amplitude in the raw signal, and averaged across all bursts (sign flipping if necessary, to make the greatest deflection positive). This method of alignment ensures that the most prominent deflection is consistently centered in the resulting potentials. This provides a clear, data-driven reference point that, importantly, is applied consistently across the different detection methods. Alternative approaches in the literature include the use of the time-frequency peak nearest to a phase minimum (Boto et al., 2022; Szul et al., 2023), which represents a valid alternative. We chose a method that does not require phase estimation within the burst. Future work could explore alternative alignment strategies.

Given the constraints on the amount of data in the mouse (Fig 3A) and LFP (Figure 3B) we simply averaged over all bursts, ignoring session or recording location. The ECoG (Figure 3C) data was different however in that we randomly chose 400 trials from each of 25 different recording sessions for one monkey, with the intent of creating an equalized dataset for this purpose. Thus, we first averaged over the trials per session and then averaged the session means, and plotted the resulting waveforms.

## Data

All data analyzed in this study was pre-existing, and in each case has been previously published with an ethics statement appropriate to that data set (Shin el al., 2017; Quilodran et al., 2008; Stoll et al., 2016). In the case of the mouse data, all experimental procedures and animal care protocols were approved by Brown University Institutional Animal Care and Use Committees and were in accordance with US National Institutes of Health guidelines. In the case of the macaque data, previously acquired in our laboratory, ethical permission was provided by the local ethical committee “Comité d’Éthique Lyonnais pour les Neurosciences Expérimentales” CELYNE, C2EA #42. ECoG data was acquired under reference C2EA42-11-11-0402-004. Animal care was in accordance with European Community Council Directives in force at the time (Directives of 1986 & 2010), and the respective implementation in French law by the Ministère de l’Agriculture et de la Forêt, with veterinary oversight from the Direction Départementale des Services Vétérinaires. All procedures were designed with reference to the recommendations of the Weatherall report, “ The use of non-human primates in research.” Laboratory authorization was provided by the “Préfet de la Région Rhône-Alpes” and the “Directeur départemental de la protection des populations” under Permit Number: #A690290402.

### Mouse

To relate each method to existing data on bursts we decided to re-analyze pre-existing data (Shin et al., 2017). This data consists of two mice, each with 5 sessions of data collection on 1 electrode of mouse barrel cortex LFP during a vibrissae deflection detection task. The sampling rate was 1000 Hz and there are approximately 200 trials per session. For both the power threshold method and cycle-by-cycle method we examined the appropriateness of the thresholds at each session, for each mouse.

#### Problem-solving task for macaque data

The macaque data are derived from two pre-existing datasets described below. In both datasets the animals are performing an explore-exploit problem-solving task first developed by Procyk & Goldman-Rakic (Procyk and Goldman-Rakic, 2006). In this task monkeys had to search an array of four possible targets by trial and error to identify the correct (rewarded) target over a series of trials. After correctly identifying the rewarded target this correct target was held for a number of trials allowing the monkey to exploit the behavior in a repetition phase. Importantly the presentation of every single trial is identical in this task – in terms of timings, visual presentation, required responses etc. The only thing that changes, other than trial outcome, between search and repetition phases is what the monkey knows about the current situation – is it searching, how well has it progressed through a search, or does it know the required response and simply need to reproduce it in the repetition phase. The analysis at the level of search and repetition phases therefore represents an analysis at the granularity of a handful of trials and provides mechanistic insight as any changes in neural dynamics between phases must be due to changes in what the monkey knows about the current trial. The task, and the search-repetition contrast, has been successful in determining a range of mechanisms within frontal neural dynamics in the contex (Procyk and Goldman-Rakic, 2006; Quilodran et al., 2008; Stoll et al., 2016; Wilson et al 2016).

In this analysis we only included trials that were from search-repetition series that once the correct target was chosen, the very same target was always chosen for the repetition phase.

### Macaque LFP

Pre-existing LFP data (Quilodran et al., 2008) was from two macaque monkeys performing an explore-exploit problem-solving task described above (Procyk and Goldman-Rakic, 2006). We included 29 (15 for monkey M, and 14 for monkey P) recording sites located in the lateral prefrontal cortex. We first low-pass filtered the data at 45 Hz and then computed the instantaneous amplitude, frequency, and phase. For our analysis we focused on 1.5 seconds of signal before the press of the target in both the search and repetition phases on the experiment. We calculated the instantaneous amplitude from 1 to 45 Hz for each trial and then rejected trials with a trial-mean amplitude greater than 2 standard deviations above the session-mean amplitude. The ratio of search to repetition was calculated at the number of trials per recording that were search trials divided by the number of trials that were repetitions. As with the mouse, in both detection methods we examined the appropriateness at each session, and for each monkey.

### Macaque ECoG

Pre-existing data from a similar problem-solving task with ECoG was also analyzed (Stoll et al., 2016). ECoG recordings are stable across many days, and we analyzed 25 different recording sessions from monkey R with one clear spectral peak in the beta band. Data were derived from ECoG recorded over the lateral prefrontal cortex, using the electrode site used for initial analyses by Stoll et al (2016). Data analyzed were from the “delay” period in that study - 1.3 seconds of the signal before the choice response. We utilized the same exact trial rejection procedure as in the LFP data above and similarly only included search-repetition series without errors in the repetition phase. Partially to equalize across sessions for later averaging and partially to cut down on computation time we randomly chose 400 trials from each session.

### Evaluation of methods in electrophysiological data

To determine the amplitude and frequency content of the bursts in each of the datasets we first computed both the instantaneous phase, frequency, and amplitude of either the raw data (LFP and ECoG) or the trial data (which we then only considered from -900 to -100 ms before the behavioral event for the burst detection for the mouse) in the beta frequency range. We then used this as a common reference for both methods, and for each detection method and each burst we calculated the mean frequency and mean amplitude over the time range of the burst. We expected the methods to pick up a slightly different burst timing, however, so small trivial changes in the mean burst characteristics could also be expected. To determine if a burst was unique to a specific method we compared each burst in a trial across methods and calculated both the proportion of temporal overlap and the relative size between each detected burst. If two bursts had an overlap greater than 0.5 and a relative size less than two times a burst then the two bursts were categorized as being common (meaning both methods picked up the same phenomenon, BoP/C). If they were not categorized as common, they were then considered unique and were assigned to whichever method detected them; either BoP or BoC.

## Supporting Information

### Simulation

We began our examination of bursts, in part, by simulating burst data to get an understanding of burst detection methods in a situation where we know the ground-truth. This was mostly done for the purposes of developing the BoC method, given the BoP method is well defined. We created 25 different simulated data sets for 4 levels of burst probability (Fig S1C). To create the twenty-five burst simulations we embedded short duration sine waves in continuous coloured noise. There were 500 trials per simulation, which included 1 second of burst-activity surrounded by a 1/2 second of noise. The sine-waves were asymmetric (rise-decay symmetry uniformly drawn between 0.25 and 0.75) with duration being uniformly drawn between 3 and 12 cycles in Fig S1. Amplitude modulation was created by multiplying the burst by an exponential function that smoothly ramped up and down. The peak amplitude could occur at 1/3, 1/2, or 2/3 after the start of the burst. Burst amplitude itself was drawn from an exponential-Gaussian function with the parameter set to 0.75, the location being set to 1e-5 and the scale set to 0.25, which is plotted in Panel D of Figure 1. In the same panel we can see that the burst frequency was drawn from a truncated normal distribution with a mean of 21, a standard deviation of 1, and being truncated between a low of 15 and a high of 29 Hz. In these simulations the main variable we manipulated was the burst probability. This was accomplished by drawing randomly from a weighted binary distribution at each time-step to determine if that time-step would start a burst or not – with there being at the least 1 cycle between bursts; the probability of the next time step was specifically set to 0.9 in the high, 0.4 in the middle, and 0.1 in the low probability manipulations. The lowest level for each of the 25 datasets was simulated 1/f noise without any intended rhythmic behavior. We then embedded periodic bursts of a varying probability of occurrence in this base data. The bursts were defined, most prominently, by their amplitude distribution, which took the shape of a log-normal distribution (Fig S1D, left), and their frequency distribution, which took a truncated normal shape (Fig S1D, right). This procedure resulted in datasets with different spectral characteristics that can be seen in Figure 1E.

**S1 Fig.**
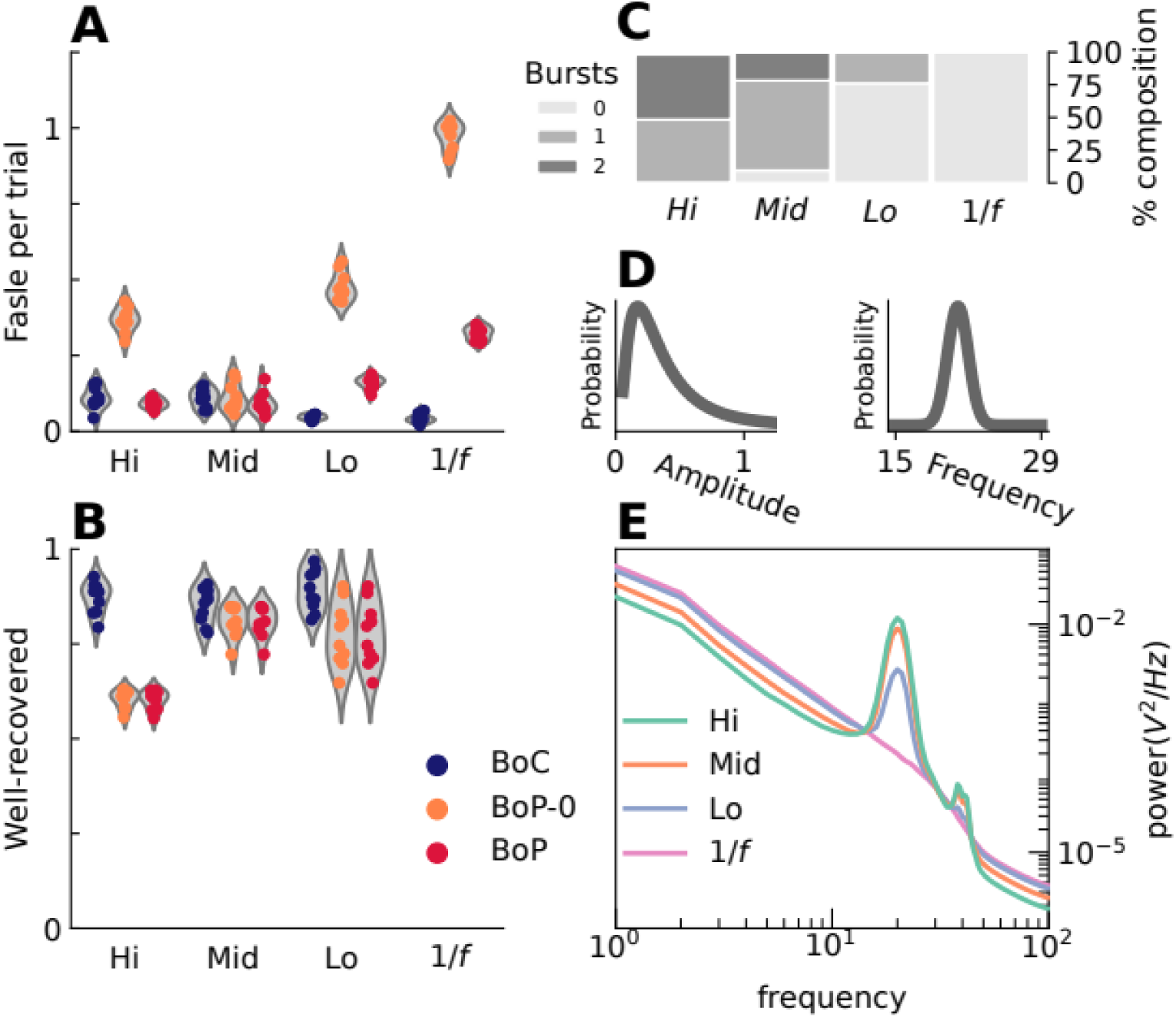
Simulation of 3-12 cycle bursts and recovery results. BoC is the Bursts of Cycle method used in the main test; BoP-0 is the Bursts of Power method without a 2-cycle constraint; and BoP is with a 2-cycle constraint. (A) The number of falsely detected bursts per trial in each of the simulation types. (B) The proportion of well-recovered bursts in each simulation. (C) The distribution of the number of bursts per simulation (100 trial s) for the four levels of burst probability. (D) Left: Burst amplitude distribution was log-normal. Right: Burst frequency distribution was a truncated normal distribution centered at 21 Hz and truncated from 15 to 29 Hz. (E) The power spectral density of each of the four levels of burst probability. **Note** that BoP and BoC target different signal features in the real electrophysiological recordings presented in the main text, and our simulations—restricted to a narrow set of burst parameters—may inadvertently favor BoC detection. Therefore, these comparisons should be considered in this context and interpreted with these limitations in mind.

We applied both the BoP-0 method and the BoC method to the simulated data sets and categorized each detected burst as being either well-recovered or not. A well-recovered burst was defined as one that shared at least half of its time points with a ground-truth burst, but was not more than twice as long. Any burst detected by a method that was not well-recovered was considered a false burst. Evaluating the performance of the BoP-0 and BoC methods using these criteria in the 3-12 cycle simulations, we found that the BoC method did not produce false bursts at high rates across all simulated data sets. In contrast, the BoP method produced false bursts at fairly high rates at both the low and high ends of the burst probability spectrum (Fig S1A). In terms of burst recovery, the BoC method was fairly consistent across all levels, while the BoC method performed poorly at the high probability level (Fig S1B). To address the issue of false bursts in the BoP-0 method, we developed a modified method that required detected bursts to be at least 2 cycles at the median frequency, and this is simply the BoP method. This simple restriction drastically reduced the number of false bursts while still recovering bursts at the same rate as the original BoP method.

## Notes

https://github.com/langfordzd/

### Competing Interest Statement

The authors have declared no competing interest.

### Summary of Updates

Rewrite of the abstract and the introduction, as well as the title change.

### Funding

- Funding for this work was provided by the Agence Nationale de la Recherche (ANR) JCJC grant “PREDYCT” to CREW: ANR-18-CE37-0016 Funding was also provided by Fondation Neurodis and by l’Institut de Neuromodulation, INM, GHU Paris Psychiatrie et Neurosciences.

